# Neural Interoceptive Processing is Modulated by Deep Brain Stimulation for Treatment Resistant Depression

**DOI:** 10.1101/2023.12.15.571885

**Authors:** Elisa Xu, Samantha Pitts, Jacob Dahill-Fuchel, Sara Scherrer, Tanya Nauvel, Jacqueline Guerra Overton, Patricio Riva-Posse, Andrea Crowell, Martijn Figee, Sankaraleengam Alagapan, Christopher Rozell, Ki Sueng Choi, Helen S. Mayberg, Allison C. Waters

**Affiliations:** Nash Family Center for Advanced Circuit Therapeutics, Icahn School of Medicine at Mount Sinai; Department of Psychiatry, Icahn School of Medicine at Mount Sinai; Department of Neurology, Icahn School of Medicine at Mount Sina; Department of Psychology, University of Arizona; Department of Psychiatry and Behavioral Sciences, Emory University School of Medicine; Department of Neuroscience, Icahn School of Medicine at Mount Sinai; School of Electrical and Computer Engineering, Georgia Institute of Technology; Department of Radiology, Icahn School of Medicine at Mount Sinai; Department of Neurosurgery, Icahn School of Medicine at Mount Sinai

## Abstract

**Background:** A critical advance in depression research is to clarify the hypothesized role of interoceptive processing in neural mechanisms of treatment efficacy. This study tests whether cortical interoceptive processing, as indexed by the heartbeat-evoked potential (HEP), is modulated by deep brain stimulation (DBS) to the subcallosal cingulate (SCC) for treatment resistant depression (TRD).

**Methods:** Eight patients with TRD were enrolled in a study of SCC DBS safety and efficacy. Electroencephalography (EEG) and symptom severity measures were sampled in a laboratory setting over the course of a six-month treatment protocol. The primary outcome measure was an EEG-derived HEP, which reflects cortical processing of heartbeat sensation. Cluster-based permutation analyses were used to test the effect of stimulation and time in treatment on the HEP. The change in signal magnitude after treatment was correlated with change in depression severity as measured by the 17-item Hamilton Depression Rating Scale.

**Results:** HEP amplitude was greater after 24 weeks of treatment (*t*(7)=-4.40, *p*=.003, *g=*-1.38, 95% Cl [-2.3, -0.42]), and this change was inversely correlated with latency of treatment response (rho = -0.75, 95% Cl [-0.95, -0.11], *p=*.03). An acute effect of DBS was also observed, but as a decrease in HEP amplitude (*t*(6) =6.66, *p*<.001, *g=*2.19, 95% Cl [0.81, 3.54]). HEP differences were most pronounced over left posterior sensors from 405-425 ms post-stimulus.

**Conclusion:** Brain-based evidence substantiates a theorized link between interoception and depression, and suggests an interoceptive contribution to the mechanism of treatment efficacy with deep brain stimulation for severe depression.

## INTRODUCTION

Deep brain stimulation (DBS) to the subcallosal cingulate (SCC) has proven effective for patients with treatment-resistant depression (TRD) (1). The immediate subjective and autonomic response to SCC DBS suggests a novel hypothesis that the treatment mechanism may involve changes in interoception (2). Interoceptive processing, or bodily sensation, is broadly implicated in the pathophysiology of depression (3,4). Understanding the effect of SCC DBS on cortical interoceptive processing in TRD, and its relationship to symptom reduction, may improve depression treatments and outcomes.

States of depression appear to modulate an electrocortical measure of interoceptive processing, the heartbeat evoked potential (HEP) (5,6). However, neural evidence of an interoceptive contribution to successful treatment for depression is lacking. The HEP reflects cortical activity that is time-locked to cardiac sensation in the chest cavity. Here, we measured the HEP longitudinally in a cohort of patients undergoing SCC DBS to test the hypothesis that successful treatment of depression will enhance cortical interoceptive processing. We also investigated immediate effects of SCC neuromodulation on the HEP as evidence that interoception is specifically relevant to the DBS treatment mechanism. This potential identification of response biomarkers is critical to advancing treatment for severe depression.

## METHODS

We enrolled 8 patients with treatment-resistant depression (Table 1) in a study testing safety and efficacy of SCC DBS (ClinicalTrials.gov #NCT01984710, #NCT04106466). Patients provided written informed consent after receiving a complete description of the study. For protocol information and study inclusion and exclusion criteria, see Supplemental Data. We assessed clinical symptom severity throughout treatment with the Hamilton Depression Rating Scale (HDRS-17) (7). Clinical response latency (time to response) was defined as the number of weeks preceding a ≥50% symptom reduction on the HDRS-17 held for a minimum of two consecutive weeks (Figure 1A). We administered the self-report multidimensional assessment of interoceptive awareness (MAIA) in 5 patients.

**Table 1.** Patients Outcome Measures.

| Subject | HDRS-17 (Baseline) | Change in HDRS-17 | Time to Response (Weeks) |
| --- | --- | --- | --- |
| 01 | 28 | -12 | 22 |
| 02 | 23 | -10.8 | 5 |
| 03 | 25 | -13.5 | 1 |
| 04 | 28 | -19 | 8 |
| 05 | 23 | -18 | 6 |
| 06 | 23 | -13.3 | 20 |
| 07 | 25 | -20.8 | 4 |
| 08 | 22 | -15.8 | 5 |

Eight patients with treatment-resistant major depressive disorder were consecutively enrolled in a study of the safety and efficacy of subcallosal cingulate deep brain stimulation for treatment-resistant depression (ClinicalTrials.gov Identifier NCT01984710, NCT04106466).
Abbreviations: HDRS-17, Hamilton Depression Rating Scale. Change in HDRS-17 calculated as the reduction in Hamilton Depression Rating Scale scores from Baseline to WK24. Time to response calculated as number of weeks until patients were at $\geq 50\%$ symptom reduction for two consecutive weeks.

**Figure 1.**
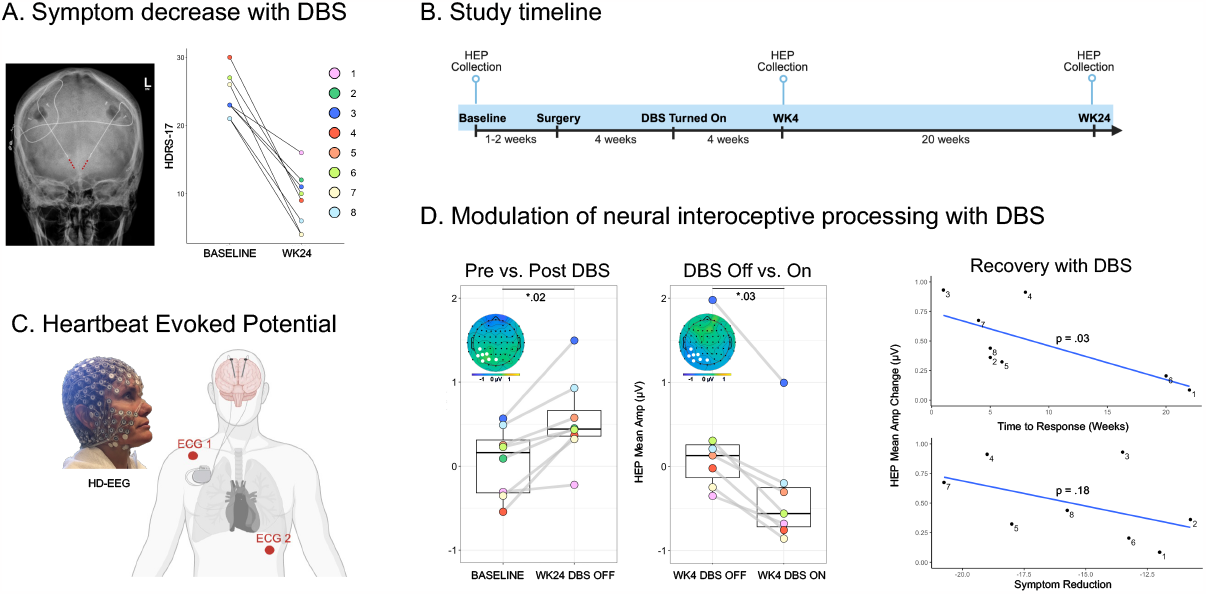
Cortical Interoceptive Processing is Modulated by Subcallosal Cingulate Deep Brain Stimulation. A. 8 patients (color coded) receiving subcallosal cingulate deep brain stimulation (DBS) for treatment resistant depression showed significant changes in depression symptom severity between baseline and 24 weeks after therapeutic DBS (WK24) measured with Hamilton Depression Rating Scale (HDRS-17). B. Study timeline highlighting 3 time points of heartbeat evoked potential (HEP) data collection: Pre-surgical Baseline, 4 weeks after therapeutic DBS turn-on (WK4), and WK24. C. HEP Data collection. Simultaneous ECG and EEG were acquired during two 3 minutes resting periods (1. DBS On, 2. DBS Off) at each data collection time point (Baseline, WK4, WK24). D. Results from cluster-based permutation analysis on the HEP between Pre vs. Post DBS (*p=*.*02)* and DBS Off vs. On *(p=*.*03)*. The topography of the HEP difference wave indicates the resulting significant difference emerged in the same location (left posterior) and time window (405-425ms). Boxplots compare each patients’ average amplitude of highlighted electrodes that consistently showed significant changes between conditions. Scatterplots show correlation between change in mean HEP amplitude and time to response (weeks) (rho = -0.75, 95% Cl, -0.95 to -0.11, *p=*.*03*), as well as HDRS-17 symptom reduction (rho = -0.52, 95% Cl, -0.29 to 0.90, *p=*.*18*).

### Procedure

Laboratory procedures were administered one to two weeks prior to surgical implantation (Baseline), 4 weeks after the date of device stimulation turn-on (WK4), and 24 weeks after the date of device stimulation turn-on (WK24). Patients were instructed to relax in a resting state with eyes closed in 3-minute blocks with simultaneous electrocardiography (ECG) and electroencephalography (EEG) recordings using a Magstim-EGI 256-channel Geodesic sensor net (1k s/s) (Figure 1C). Experimental conditions (DBS On; DBS Off) were used to evaluate the acute effect of stimulation after 4 and 24 weeks of treatment. In these acute DBS On conditions, patients received the same clinically defined, therapeutic parameters used for chronic DBS: 130 Hz; 90 µs pulse width, 3-5 V. See Figure 1B for study timelime; for more details about EEG acquisition, see Supplemental Data.

### Preprocessing and extraction of the Heartbeat Evoked Potential

Preprocessing of the EEG data and extraction of the HEP were completed in NetStation software (Magstim-EGI, Eugene, OR) and Matlab v2022b (8). EEG data were re-referenced to the average reference and downsampled to 70 channels. EEG was then bandpass filtered between Hz and 30 Hz to account for low frequency noise and the DBS stimulation artifact.

Non-neural artifact rejection was performed using Matlab v2022b with the eeglab2022_1 toolbox (9). Independent component analysis was used to reject the cardiac field artifact (CFA), which represents an important potential confound for investigations of the HEP (10). For a more detailed discussion of artifact rejection steps, including the assessment of the CFA, see Supplemental Data. A custom algorithm was designed in Matlab v2022b to detect the R-peak of the ECGs. Continuous EEG was epoched around the ECG’s R-peak to extract the heartbeat evoked potential. Epochs had a duration of 850 ms, starting 200 ms before the R-peak and ending 650 ms after. A 150 ms time window preceding the R-peak onset was used to baseline correct the HEP.

### Data analysis

A comparison of psychometric scores pre/post DBS, as well as the correlation of change scores with neural measures, were conducted in RStudio (11). The effect of time in treatment on HEP magnitude was assessed with stimulation off, to maintain methodological consistency with the presurgical baseline: Baseline versus WK24. The acute effect of stimulation on HEP magnitude was assessed at WK4 and WK24 of treatment, separately: DBS ON vs. DBS OFF. Statistical contrasts were made in Matlab v2022b with a mass-univariate analysis at all sample time points and electrode locations between 400-500 ms after the ECG R-peak, consistent with previously reported methods (10,12). Detailed parameters of the permutation analysis are provided in Supplemental Data.

## RESULTS

Sample characteristics, provided in Table 1, show HRDS-17 scores at baseline (Mean= 25, SD = 2.3), and week 24 (Mean = 9.1, SD = 4) with a treatment response in 6 of 8 patients. No statistical difference was detected (Baseline vs. WK24) in MAIA total and subscale scores (*p>0*.*05*).

Cluster-based permutation analyses revealed significant differences between conditions in both Baseline versus WK24 DBS Off (*p* =.02) and WK4 DBS On versus WK4 DBS Off (*p* = .03) contrasts in the same left posterior location and time window (405-420 ms) (Figure 1D). There was a significant difference in the HEP amplitude between Baseline and WK24 DBS Off, showing a more positive HEP amplitude for the WK24 DBS Off condition than Baseline (cluster electrodes, *t*(7)=-4.40, *p*=.003, *g=*-1.38, 95% Cl [-2.3, -0.42]). Additionally, there was a significant difference in the HEP amplitude between WK4 DBS On and WK4 DBS Off, showing a more negative HEP amplitude for the WK4 DBS On condition than WK4 DBS Off (cluster electrodes, *t*(6) =6.66, *p*<.001, *g=*2.19, 95% Cl [0.81, 3.54]). WK24 DBS On vs. WK24 DBS Off revealed no significant effect of acute DBS (*p= >*.*37*). More detailed permutation results are discussed in Supplement 1.

Change in HEP amplitude over 24 weeks of treatment was inversely correlated with latency of treatment response (rho = -0.75, 95% Cl [-0.95, -0.11], *p=*.03) (Figure 1D). A similar relationship was observed with HDRS-17 scores, but below threshold for statistical significance (rho = 0.52, 95% Cl [-0.29, 0.90], *p=*.18) (Figure 1D). Change in HEP amplitude was not correlated with change in MAIA.

## DISCUSSION

We examined the effect of SCC DBS treatment for severe depression on neural interoceptive processing. All patients showed symptom improvement after 6 months of DBS as well as an enhancement in neural interoceptive processing. Individuals with the most rapid treatment response showed the largest increase in the HEP cortical signal, suggesting a relationship between the antidepressant response trajectory and changes in neural interoceptive processing. In contrast to the enhanced interoceptive signal after chronic DBS, acute DBS in the laboratory caused an immediate suppression of the interoceptive signal. This effect was observed early in treatment and converges with previous evidence of an inversion in SCC DBS treatment mechanisms between acute and chronic phases of the treatment response (13).

Without a behavioral measurement of interoceptive experience, our interpretation assumes that the heartbeat evoked potential (HEP) indexes cardiac interoceptive processing. This is supported by evidence that the HEP localizes to core interoceptive regions and is modulated by interoceptive attention, arousal and self-reference (14). Notably, subjective self-report measures of interoceptive awareness showed no change in this sample. However, self-report measures may be insufficient to establish HEP as a definitive measure of interoception, reflecting dissociable dimensions of interoceptive experience (15). Meanwhile, the consistency of our results to reported HEP topography and latency lends confidence to our conclusion, particularly given the rigorous statistical constraint applied to a small sample. Similarly, correlative outcomes are speculative in a small sample, but are consistent with the hypothesized relationship between symptoms of depression and cortical interoceptive processing.

A key interoceptive contribution to mood disorders and the SCC DBS treatment mechanism, specifically, has been proposed but not demonstrated objectively (2). We provide this critical evidence at a moment when clarifying the SCC DBS treatment mechanism is essential to further development of this lifesaving treatment. Intervening on aberrant interoceptive neural processes may be key to enhancing DBS and other approaches to treatment resistant depression.

## Acknowledgments

We sincerely thank the patients and their families for their trust, commitment and collaboration that enabled this study. DBS devices used in this research were donated by Medtronic (Minneapolis, MN).

## Previous presentations

1. Poster presentation at Society of Biological Psychiatry 2023 Annual Meeting (April, 2023)
  a. Xu E, Pitts S, Dahill-Fuchel J, Scherrer S, Overton J, Nauvel T, Posse PR, Crowell A, Figee M, Gowatsky J, Alagapan S. 280. Enhancement of Neural Interoceptive Processing Observed in Responders to Deep Brain Stimulation for Treatment Resistant Depression. Biological Psychiatry. 2023 May 1;93(9):S206-7.
2. Oral presentation at Society for Psychophysiological Research 2023 Annual Meeting (September, 2023)

