## Supplemental Material for "Neural Interoceptive Processing is Modulated by Deep Brain Stimulation for Treatment Resistant Depression"

**Methods.**
**Results.**
**Figure 1.** Rastor Plot of HEP Group Differences: Baseline Compared with WK24 HEP
**Figure 2.** Rastor Plot of HEP Group Differences: WK4 DBS On Compared with WK4 DBS Off
**Figure 3.** Rastor Plot of HEP Group Differences in Frontal Electrodes: WK4 DBS On Compared with WK4 DBS Off
**References.**

This supplementary material has been provided by the authors to give readers additional information about their work.

### Methods

#### Study Population

The protocol adhered to the Declaration of Helsinki and was approved by the Institutional Review Boards of Emory University and the Icahn School of Medicine at Mount Sinai, as well as the US Food and Drug Administration under a physician-sponsored Investigational Device Exemption (IDE G130107) and is monitored by the Emory University Department of Psychiatry and Behavioral Sciences Data and Safety Monitoring Board. All participants signed an informed consent to participate; all patients continue in the ongoing longitudinal study.

Eight subjects with treatment-resistant major depressive disorder were consecutively enrolled in a study of the safety and efficacy of SCC-DBS for treatment-resistant depression (ClinicalTrials.gov Identifier NCT01984710). The study inclusion and exclusion criteria were identical to those previously published (1,2). Participant race, ethnicity, sex, and gender were self-reported in the context of a structured clinical interview. In brief, patients had a depressive episode of at least one year duration, a Hamilton Depression Rating Scale (HDRS-17) (3) severity score of 20 or higher; trials of at least 4 antidepressant treatments (including electroconvulsive therapy) without improvement; no significant psychiatric or medical comorbidities; and significant functional impairment with a Global Assessment of Function score (4) of less than 50 (range, 1–100, with higher scores indicating better function). Clinical symptom severity was assessed by an independent rater using the HDRS-17 during weekly visits to the laboratory among other behavioral scales. Following established criteria, a decrease in HDRS-17 scores greater than 50% of the presurgical average was set as the threshold for 'response'.

#### Psychometric Assessment

The multidimensional assessment of interoceptive awareness (MAIA) (5) was administered with a subset of patients (n=5) at baseline and WK24. These patients include 01, 02, 03, 04, and 05. MAIA is a self-report questionnaire that consists of 32 items assessing 8 different dimensions of interoception: "(1) Noticing: the awareness of uncomfortable, comfortable and neutral body sensations; (2) Not-Distracting: the tendency to ignore or distract oneself from sensations of pain or discomfort; (3) Not-Worrying: emotional distress or worry with sensations of pain or discomfort; (4) Attention Regulation: the ability to sustain and control attention to body sensation; (5) Emotional Awareness: the awareness of the connection between body sensations and emotional states; (6) Self-Regulation: the ability to regulate psychological distress by attention to body sensations; (7) Body Listening: actively listening to the body for insight and (8) Trusting: experiencing one's body as safe and trustworthy." Each item of the different subscales is a Likert scale with six levels ranging from 0 (never) to 5 (always).

#### EEG recording procedure:

Hardware and software used to acquire and process EEG were products of Electrical Geodesics Inc., (Eugene, OR). Patients were fitted with a 256-channel Hydrocel Geodesic Sensor Net and seated in a climate controlled room. A chin rest was used to reduce motion artifacts. Electrode impedance were maintained below 50 kOhm. Recordings were made at a rate of 1,000 samples per second through a NetAmps 400 ampilfier using NetStation software. Patients were instructed to relax and let their mind wander.

#### Preprocessing and extraction of the Heartbeat Evoked Potential:

Preprocessing of the EEG data and extraction of the HEP were completed in NetStation software (Electrical Geodesics, Eugene, OR) and Matlab v2022a (Mathworks, Natick, MA). A custom detection algorithm was designed in Matlab v2022a to detect the R-peak of the ECGs. The EEG montage was down-sampled to 70 channels and re-referenced to the average reference using a polar average reference effect (PARE) correction to estimate the zero surface potential integral (6) on NetStation software. EEGs were then bandpass filtered between 0.1 Hz and 30 Hz to account for low frequency noise and the DBS stimulation artifact.

Further data preprocessing including artifact rejection and detection was performed using MATLAB_R2022b with the eeglab2022_1 toolbox. Artifact Subspace Reconstruction (ASR) was applied to reduce high-amplitude artifacts (7). The ASR parameter of 30 was chosen to balance between removing non-brain signals and retaining brain activities. Continuous EEG was epoched around the EKG's R peak to extract the heartbeat evoked potential (HEP). Epochs had a duration of 850 ms, starting 200 ms before the R peak and ending 650 ms after. The epoched data was subsequently manually inspected for further ocular, muscular, and movement artifact detection and rejection.

**Assessment of Cardiac Field Artifact**

The cardiac field artifact (CFA) represents an important potential confound for investigations of the HEP, with no universally agreed upon solution. One generally powerful method for artifact correction of EEG data is independent component analysis (ICA) (8). The ICA identified the CFA as independent components, which we removed following the examples provided by previous research (9,10,11,12). As an additional check, we also analyzed the ECG trace mimicking the permutation analysis procedure followed in the HEP analysis (see below).

**HEP Analysis and Statistics:**

Event-related potential analysis was performed by averaging the EEG epochs for Baseline vs. WK24 DBS Off, WK4 DBS On vs. WK4 DBS Off, and WK24 DBS On vs. WK24 DBS Off. Segments of EEG were visually inspected to identify and interpolate channels with excessive noise. An average of 6.5 channels (SD= 3.69) were interpolated using spherical-spline interpolation, using EEGLAB toolbox. The data was then re-referenced to the average reference.

In light of methodological variations in HEP amplitude measurement in the literature, specifically pertaining time window and electrode selection for analysis, we conducted a mass-univariate analysis at every time point and electrode location with a spectrum of 400-500 ms after the R peak. This approach and time window was selected based on a systematic review and meta-analysis on HEP and interoception.

To detect reliable differences between the HEP at Baseline vs. WK24 DBS Off, the HEP at WK4 DBS On vs. WK4 DBS Off, and the HEP at WK24 DBS On vs. WK24 DBS Off, the HEPs from these conditions were respectively submitted to a repeated measures, two-tailed cluster-based permutation test based on the cluster mass statistic[13] using a family-wise alpha level of 0.05. The Mass Univariate ERP Toolbox was utilized for this examination. Our focus was on 100 time points spanning from 400 to 500 ms, observed across 70 scalp electrodes, resulting in 7000 total comparisons. For each comparison, we performed repeated measures t-tests using both the original data and 2500 random within-participant permutations of the data. In each permutation, t-scores corresponding to uncorrected p-values of 0.05 or lower were grouped into clusters, with neighboring t-scores forming a cluster. Electrodes within a proximity of approximately 4.99 cm were considered spatial neighbors, and adjacent time points were deemed temporal neighbors. The cumulative t-score within a cluster represented its "mass." Among the 2501 test sets, the most extreme cluster mass was recorded from each and used to estimate the null hypothesis distribution (i.e., no difference between conditions). The observed data's permutation cluster mass percentile ranking was employed to derive the p-value for each cluster. T-scores not part of a cluster were assigned a p-value of 1.

We chose this permutation test analysis over the more traditional mean amplitude ANOVAs due to its superior spatial and temporal resolution, while still moderately controlling the family-wise alpha level (accounting for the numerous comparisons). Additionally, we opted for the cluster mass statistic in this permutation test due to its favorable power characteristics in capturing widely distributed ERP effects like the P300 (14,15). The decision to employ 2500 permutations was guided by the fact that it exceeds the number recommended by Manly (1997) for a family-wise alpha level of 0.05 (16), enhancing the robustness of our analysis.

### Results

#### Null Effect of Cardiac Field Artifact on Experimental Contrasts

The results of the cluster-based permutation test on the ECG did not reveal any cluster of significant interactions at $P < 0.05$, corrected for multiple comparisons. In the absence of any differences in cardiac activity between conditions, it is safe to assume that the CFA is constant across and will therefore not affect the contrast between Baseline vs. WK24 DBS Off, nor the contrast between WK4 DBS On vs. WK4 DBS Off.

#### Null Effects of Data Attrition on Experimental Contrasts

For the change over time (Baseline vs. WK24 DBS Off) contrast, an average of 58% of epochs were rejected for Baseline, leaving an average of 286.75 (SD= 63.95032) of epochs across the 8 participants. An average of 31.8% of epochs were rejected for WK24 DBS Off, leaving an average of 198.125 (SD= 50.54683) of epochs per participants. We found no significant difference in number of epochs between Baseline and WK24 DBS Off (Wilcoxon Signed-Rank Test $z = 1.82$, $p = 0.080$). For the effect of acute stimulation (WK4 DBS On vs. WK4 DBS Off) contrast, patient 802 was excluded due to an excessively noisy EEG recording despite data cleaning procedures. An average of 15.9% of epochs were rejected for WK4 DBS On, leaving an average of 220 (SD= 47.35) of epochs across the remaining 7 participants. An average of 23.2% of epochs were rejected for WK4 DBS Off, leaving an average of 199.2857 (SD= 51.82572) of epochs per participants. We found no significant difference in number of epochs between C4 DBS On and C4 DBS Off (Wilcoxon Signed-Rank Test $z =1.18$, $p = 0.272$).

#### Permutation tests

*Baseline vs. WK24 DBS Off*

Using a cluster-based permutation approach, we tested for changes in the HEP as a function of treatment change over time with. We found a significant difference in the HEP amplitude between Baseline and WK24 DBS Off, showing a more positive HEP amplitude for the WK24 DBS Off condition than Baseline, where midline-posterior electrodes consistently show the significant effect. Specifically, the cluster ranged from 400 to 500 ms and the spatial extent was between left lateralized posterior electrodes and right lateralized posterior electrodes. (**Figure 1**)

*WK4 DBS On vs. WK4 Dbs Off*

Using the same approach, we also tested for changes in the HEP as a function of acute DBS at WK4. We found a significant difference in the HEP amplitude between WK4 DBS On and WK4 DBS Off, showing a more negative HEP amplitude for the WK4 DBS On condition than WK4 DBS Off, where midline-posterior electrodes consistently show the significant effect. Specifically, the cluster ranged from 400 to 473 ms and the spatial extent was between left lateralized posterior electrodes and right lateralized posterior electrodes. (**Figure 2)**

*WK24 DBS On vs. WK24 DBS Off*

We tested for changes in the HEP as a function of acute DBS at WK24. We did not find a significant difference in the HEP amplitude between WK24 DBS On and WK4 DB2S Off.

#### Exploratory/Supplemental Analyses

To select an appropriate montage for further analyses with behavioral measures, electrodes that consistently showed significant changes across both contrasts in the aforementioned location and time window were chosen (CP5, P7, P5, P3, PO7, PO3, POz).

#### WK4 DBS On vs. DBS Off Frontal Electrodes

Prior literature implicated the HEP in frontal midline electrodes. By only including frontal-midline electrodes in the permutation test comparing DBS On vs. DBS Off at WK4 while keeping the time window of 400 – 500 ms constant, we were able to find a significant difference in the HEP amplitude, showing a more positive HEP amplitude for the WK4 DBS On condition than WK4 DBS Off, where frontal midline electrodes consistently show the significant effect. Specifically, the cluster ranged from 405 to 464 ms and the spatial extent was between left lateralized frontal electrodes and frontal central electrodes (p=.05). However, this effect does not hold without excluding the posterior electrodes from the permutation test. **(Figure 3)**

**WK24 DBS On vs. DBS Off**

While a cluster-based permutation approach at WK24 (WK24 DBSOn minus WK24 DBSOff) revealed no significant effect of acute DBS, DBS On may differ from DBS Off at WK24 as our results may have been underpowered (p=0.36). If we apply the left posterior electrode montage ascertained from previous contrasts and test for differences between conditions in the 405-420 ms time window, we observe a similar directional effect of DBSOn as in WK4 in every patient except for Patient 1.

**Correlation between Change in HEP Over Time and Change in MAIA Trusting Scale**

In the subset of patients (n=5) who completed the MAIA questionnaire at baseline and WK24, we tested if the change in HEP amplitude over 24 weeks of treatment is related to changes in subjective interoceptive awareness. Subscales of MAIA change over time did not significantly correlate with change in HEP over time. However, change in the MAIA subscale “Trusting” tracked with symptom change, such that greater change in HEP amplitude may be associated with a greater increase in “Trusting” scores (*rho* = 0.7, p =n.s.).

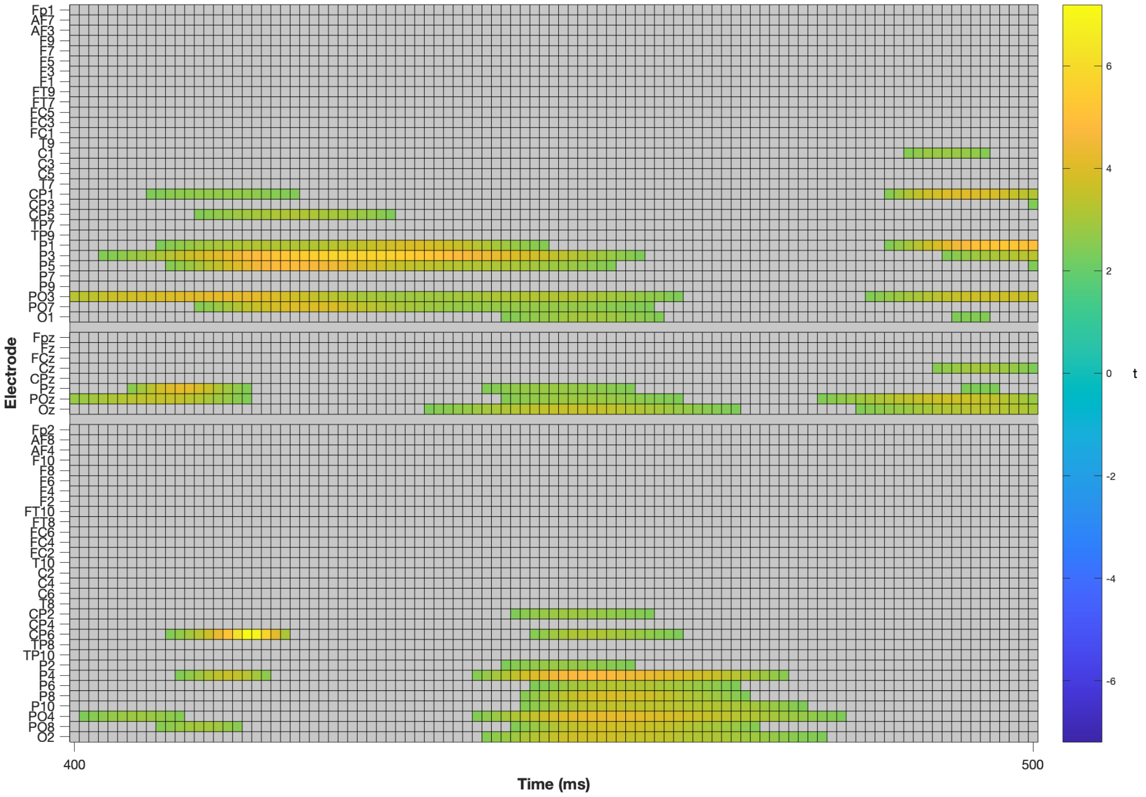


**Figure 1.** Rastor Plot of HEP Group Differences: Baseline Compared with WK24 HEP
Raster plot demonstrating significant Baseline versus WK24 group differences for HEP. The raster depiction showcases significant differences observed within the experimental group, as detected by the permutation test utilizing the cluster mass statistic. Each colored square of electrode and time point denotes a significant p-value ($p < .05$). Areas with no significant impact are represented by grey rectangles. It's important to acknowledge that the arrangement of electrodes follows a topographical arrangement along the y-axis. Electrodes situated on the left and right sides of the head are respectively grouped at the top and bottom of the illustration. Meanwhile, midline electrodes are positioned in the middle. Within these three groupings, the y-axis orientation from top to bottom corresponds to the anterior-to-posterior progression across the scalp.

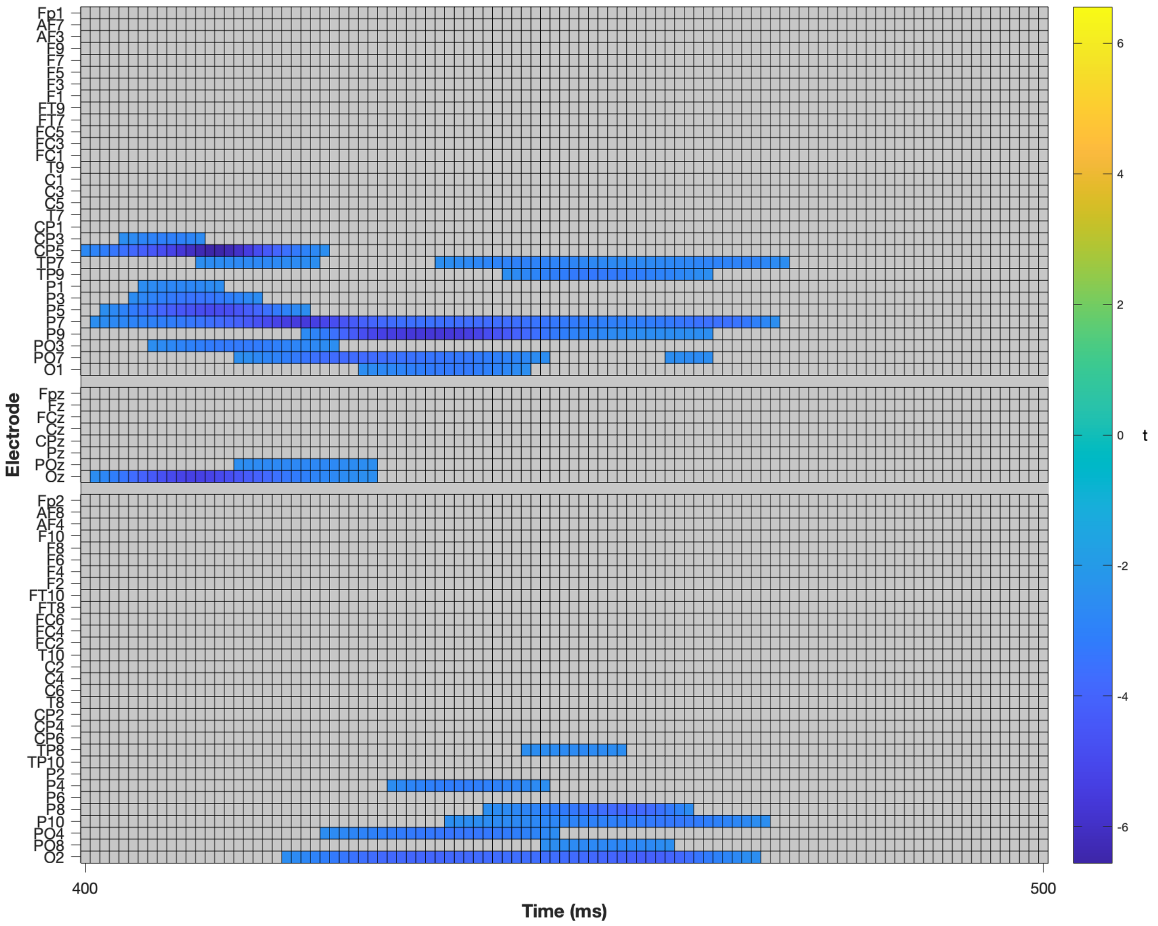


**Figure 2.** Rastor Plot of HEP Group Differences: WK4 DBS On Compared with WK4 DBS Off
Raster diagram illustrating significant WK4 DBS On versus WK4 DBS Off group differences for HEP. The raster depiction is organized as in Figure 1 above.

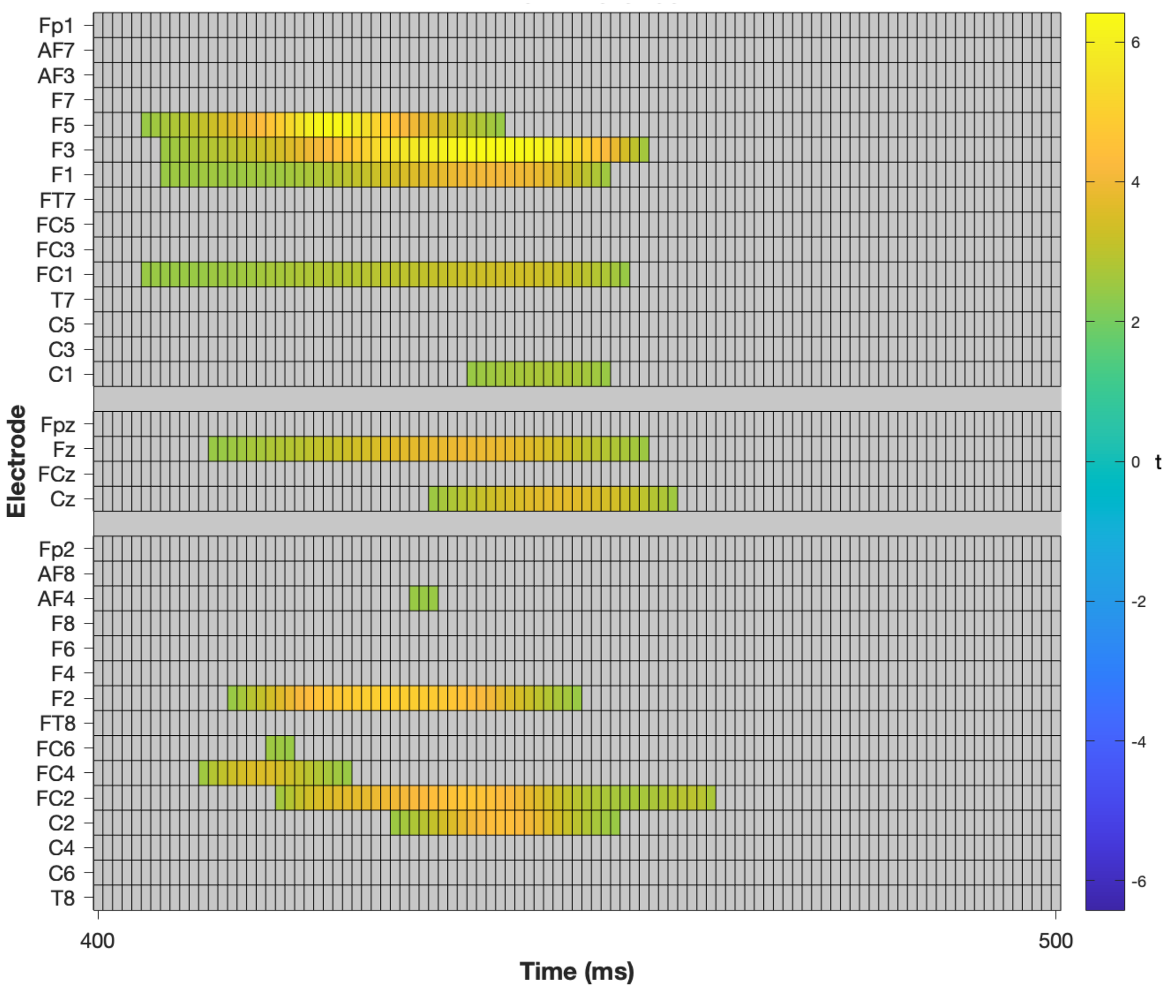


**Figure 3.** Rastor Plot of HEP Group Differences in Frontal Electrodes: WK4 DBS On Compared with WK4 DBS Off

Raster diagram illustrating significant WK4 DBSOn versus WK4 DBSOff group differences for HEP with permutation restricted to frontal midline electrodes. The raster depiction is organized as in Figure 1 above.

### Supplemental Data References


1. Holtzheimer PE, Kelley ME, Gross RE, Filkowski MM, Garlow SJ, Barrocas A, ... Chismar R. Subcallosal cingulate deep brain stimulation for treatment-resistant unipolar and bipolar depression. Arch Gen Psychiatry. 2012;69(2):150-158.
2. Riva-Posse P, Choi K, Holtzheimer PE, Crowell AL, Garlow SJ, Rajendra JK, ... Mayberg HS. A connectomic approach for subcallosal cingulate deep brain stimulation surgery: prospective targeting in treatment-resistant depression. Mol Psychiatry. 2017.
3. Hamilton M. A rating scale for depression. J Neurol Neurosurg Psychiatry. 1960;23(1):56.
4. Hall RC. Global assessment of functioning: a modified scale. Psychosomatics. 1995;36(3):267-275. (Junghöfer, Elbert, Tucker, & Braun, 1999)
5. Mehling WE, Price C, Daubenmier JJ, Acree M, Bartmess E, Stewart A. The multidimensional assessment of interoceptive awareness (MAIA). PLoS One. 2012;7(11):e48230.
6. Junghöfer M, Elbert T, Tucker DM, Braun C. The polar average reference effect: A bias in estimating the head surface integral in EEG recording. Clin Neurophysiol. 1999;110(6):1149–1155.
7. Chang CY, Hsu SH, Pion-Tonachini L, Jung TP. Evaluation of artifact subspace reconstruction for automatic artifact components removal in multi-channel EEG recordings. IEEE Trans Biomed Eng. 2019;67(4):1114-1121. (Debener, Thorne, Schneider, & Viola, 2010; Hoffmann & Falkenstein, 2008)
8. Devuyst S, Dutoit T, Stenuit P, Kerkhofs M, Stanus E. Cancelling ECG artifacts in EEG using a modified independent component analysis approach. EURASIP J Adv Signal Process. 2008;2008:1-13. (Hoffmann & Falkenstein, 2008)
9. Villena-Gonzalez M, Rojas-Thomas F, Morales-Torres R, López V. Autonomous sensory meridian response is associated with a larger heartbeat-evoked potential amplitude without differences in interoceptive awareness. Psychophysiology. 2023:e14277.
10. Mai S, Wong CK, Georgiou E, Pollatos O. Interoception is associated with heartbeat-evoked brain potentials (HEPs) in adolescents. Biol Psychol. 2018;137:24–33.
11. Terhaar J, Viola FC, Bär KJ, Debener S. Heartbeat evoked potentials mirror altered body perception in depressed patients. Clin Neurophysiol. 2012;123(10):1950–1957.
12. Luft CDB, Bhattacharya J. Aroused with heart: Modulation of heartbeat evoked potential by arousal induction and its oscillatory correlates. Sci Rep. 2015;5(1):15717. (Bullmore et al., 1999)
13. Bullmore ET, Suckling J, Overmeyer S, Rabe-Hesketh S, Taylor E, Brammer MJ. Global, voxel, and cluster tests, by theory and permutation, for a difference between two groups of structural MR images of the brain. IEEE Trans Med Imaging. 1999;18(1):32-42.
14. Groppe DM, Urbach TP, Kutas M. Mass univariate analysis of event-related brain potentials/fields II: Simulation studies. Psychophysiology.
15. Maris E, Oostenveld R. Nonparametric statistical testing of EEG- and MEG-data. J Neurosci Methods. 2007;164(1):177-190.
16. Manly BFJ. Randomization, Bootstrap, and Monte Carlo Methods in Biology (2nd ed.). London: Chapman & Hall; 1997.
